# Persistent homology centrality improves link prediction performance in Pubmed co-occurrence networks

**DOI:** 10.1101/2024.03.19.585668

**Authors:** Chase Alan Brown, Jonathan D. Wren

## Abstract

This paper provides a novel approach to understanding the nature of innovation and scientific progress by analyzing large-scale datasets of scientific literature. A new measure of novelty potential or disruptiveness for a set of scientific entities is proposed, based in the mathematical formalism of algebraic topology via a method called *persistent homology*. In this framework, understanding where academic ideas depart from the existing body of knowledge to *fill knowledge gaps* is key to scoring a set of entities and their potential for filling future knowledge gaps. This framework is motivated by the assumption that scientific discovery has underlying regularities that can be modeled and predicted.

Our method uses a *filtration*, which is a type of ranking of hypergraph components along a chosen parameter. In this work two different axes are used, which constructs a growing grid of sub-hypergraphs. The axes of time (scientific knowledge evolution) and normalized point-wise mutual information (network structure) affords the ability to succinctly represent the entire dynamic structure of the scientific literature network. We then find that using very simple and interpretable measures of centrality derived from this crude *bifiltration* or *vineyard* affords the ability to predict links within the dynamic scientific network.

While several different methods of link prediction have been proposed in the past, the method presented here *extends* the notion of link prediction to a higher dimension, as the boundary of the knowledge gap may be more than just 0-dimensional nodes.

The system presented here not only suggests a mathematical basis, consistent with observations in cognitive neurosciences regarding early childhood language acquisition, but additionally provides useful applications for the scientific community in predicting and ranking hypothesis for scientific discovery.

## 1 Introduction

### Algebraic Topology in Neuroscience, Machine Learning, and Cognitive Sciences

The field of algebraic topology (AT) has produced remarkable new discoveries in both neuroscience and machine learning, from the biological observation of toroidal activity in grid cells, to the cyclical arrangement of embeddings in the well known “grokking” artificial neural network (ANN) mechanistic interpretability experiment. ^1–3^

Furthermore, this branch of mathematics has been used as a data analysis tool-set to provide an observation that childhood semantic growth networks (CSGNs) form and fill ‘knowledge gaps’ (`k`_gaps_) in a very predictable and structured pattern - via a nice mathematical tool from AT called *persistent homology* (PH).^4^ The `k`_gap_ definition used within our work is the same as that used in this CSGN study: a representative of a generator Γ found using PH. That is, a set of vertices and edges from a knowledge graph, connected together to form a ‘boundary’ (closed loop) of dimension *k*. This work continues with this same algebraic topology-based methodology, applying it to the prediction of scientific discovery in literature. To this end, we present a system that applies PH to a literature co-occurrence network^5^ of biological and cancer hallmark entities, which can generate and rank hypotheses for scientific discovery.

### Scientific discovery operationalization with interpretable methods

This research aims to integrate the broad spectrum of philosophical perspectives and academic disciplines that study the scientific innovation and discovery process. We draw upon several foundational fields in this area, including classical philosophy, literature-based discovery, cognitive neuroscience, and topological data analysis. Classical works have historically given insight into descriptions of the mechanisms by which discovery occurs and the nature of its evolution.^6^ However, with the recent progress in language modeling, which will continue to help facilitate knowledge-based work, our research here looks to address one of the key problems lacking in large language model (LLM) capabilities at the time of writing - producing novel, insightful, and highly plausible hypotheses in an *interpretable* and *operationalized* manner. Consequently, a primary interest of this work is to bridge the divide between the qualitative philosophies of epistemology, and the quantitative sciences characterized by robust predictive power.

While modern LLMs at present have inherent constraints on autonomously generating precise goals and hypotheses without explicit instruction, one objective for this study is to provide a novel methodology that mitigates this deficiency, and provides starting hypotheses for researchers which utilize the full background of scientific literature. Additionally, the system proposed not only directs the construction of hypotheses, but also empowers researchers to more quantitatively assess their position or grant ideas within the full background of scholarly publications.

### Utilizing hints from cognitive neurosciences

We aim to provide a new perspective on the problem of how language can be properly represented through the lens of *topological data analysis* (TDA) and *persistent homology*, as they have been shown to be a powerful tool in cognitive neuroscience (in the study of early CSGNs^4^), the study of the brain’s connectome (both macroscopically and microscopically), as well as in the study of ANNs.

## 2 Previous Works

Studying the nature of discovery is not entirely new, and has been framed over the years in various ways by Thomas Kuhn^6^, Karl Popper^7^, Imre Lakatos^8^, and many others. The study of the field of discovery is founded upon the premise that the process of discovery is not a random process, but rather a process that is governed by underlying principles.

In this paper, we work within this assumption by improving a standard model of link prediction in growing knowledge graphs, while using a methodology which has been shown to be effective in predicting the growth of children’s semantic networks. Therefore, our motivation is to test whether the same principles from cognitive neuroscience that govern children’s semantic growth networks (CSGNs) and language development also govern the growth of scientific knowledge.

### 2.1 Childhood Semantic Growth Networks

In previous work from Ann Sizemore^4^, it has been shown that children’s semantic growth networks can be modeled and predicted using TDA. This work has shown that the formation and filling of `k`_gaps_ in children’s semantic growth networks is how the process of learning occurs at an early age (e.g. mommy, daddy, bottle, ball, etc.). This work has also shown that the filling of `k`_gaps_ is a universal process, independent of cultural and linguistic differences in children across the world.

This work is an extension of a large body of papers in a field that studies early childhood development. One prominent and fundamental work in this field was establishing the WordBank dataset^9^, which is built upon the MacArthur-Bates Communicative Development Inventory (MB-CDI). The MB-CDI is a type of questionnaire/survey that parents can complete to report early language information about their children. One of the first reports using the MB-CDI consisted of 1,803 children, ages 8-30 months^10^ and determined some of the regularities of early language learning. As of the time of writing, the WordBank dataset contains data from ∼93k children (105,290 CDI administrations) across 42 languages. Many studies in this field continue to use this dataset to show structure within the dynamics of CSGNs^11,12^ while making use of these foundational datasets, which can be used to create the edges in the CSGNs, via connecting ‘features’ of the words for showing `k`_gap_ formation.^13^

In the CSGN paper from Sizemore^4^, persistent homology was key in determining that the evolution of semantic networks have regularities in their dynamics which can be observed in the barcodes, persistence diagrams, or Betti curves of the data. In our work, we use this observation in order to improve link prediction in growing Pubmed Co-occurrence networks.

### 2.2 Wikipedia growth networks

Extending upon the work from CSGNs, it has been observed that the growth of Wikipedia articles can be modeled and predicted using this same TDA framework.^14^ The premise of the papers on Wikipedia growth networks closely related to the work presented in this paper; though with key differences in methodology and scope. First, the data used in this paper is from the scientific literature, and not from the aggregated and filtered knowledge base of Wikipedia.

Wikipedia is a much smaller subset of the data considered in this work, does not include the full complexity of ideas available to scientists in journals, and does not update at the same frequency to include the plethora of information presented to researchers on a daily basis.

Our dataset, based on all articles in Pubmed, is a more accurate representation of the full scope of scientific knowledge, and therefore has the potential to be more practically useful for predicting discoveries. Additionally, the representation used within the Wikipedia growth networks is a directed graph structure constructed from the hyperlink structure of Wikipedia articles, and not a co-occurrence network from applying information retrieval techniques, such as named entity recognition (NER) and entity-linking (EL) to the language within the abstracts directly. This difference in representation is important, as hyperlinks are far more sparse and do not always encode the same information as all entities within the text.

These limitations in data update frequency and completeness motivate our work towards a tool that can predict scientific discovery in a more accurate and timely manner that will be more useful for researchers in a practical way.

## 3 Methods

### 3.1 Data

The dataset used in this study is *LION-LBD*^5^, which is a set of graphs decorated with various node and edge data, and is freely and publicly available at https://lbd.lionproject.net/downloads ^15^. The dataset is derived from a collection of scientific articles from Pubmed, and the graphs are constructed by connecting biologically related entities (genes, drugs, hallmarks of cancer, etc.) that co-occur in the same article abstract. The weight of each entity pair is calculated in many ways, including normalized point-wise mutual information (NPMI, or ν), which is the most used metric for the analyses within this paper.

The dataset consists of 10 sub-graphs, each constructed to probe a different concept space, which were further split by the *LION* group into 5 “landmark” discovery sets for modern (∼2017) era and 5 “Swanson linking” (historical) datasets. The datasets are referred in this work as 𝒟_𝓌_ ∈ **𝒟** where *w* is each concept space. These datasets are enumerated in Table 1.^15^ Each dataset is a *dynamic* graph, decorated with additional information. Recall that a simple graph can be defined as 𝒢 = (*V, E*) and {*e* ∈ *E* | *e* = (*v*_*i*_, *v*_*j*_) ∧ *v*_*i*_, *v*_*j*_ ∈ *V* }, where *V* is the set of vertices (nodes), and *E* the set of edges.

The datasets for this work have some other information: 𝒢 = (*V, E*, Φ_*ν*_, Φ_*e*_). A graph at a particular time can be extracted with the relevant fields to construct a graph from the dataset with a simple function *f*_extract_ : (𝒟_𝓌_, *t*) → 𝒢_*w,t*_ which will then produce a graph ‘decorated’ with additional information at the desired time.

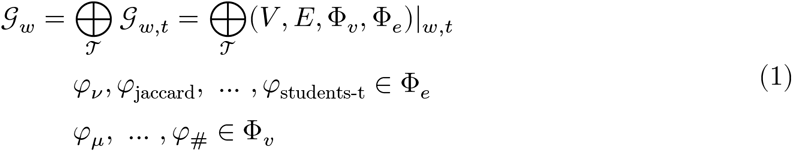

Where *w* is the sub-dataset from LION-LBD, *t* is the time or time-slice from the ordered set of times 𝒯, *φ*_*ν*_(*v*_*i*_, *v*_*j*_) is the NPMI function, each edge *V* is the set of nodes (vertices), and *E* is the set of edges . Additionally, the graphs are strictly growing in time:

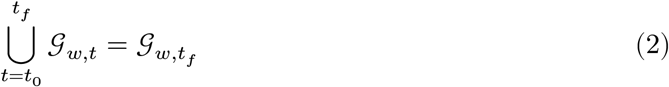

### 3.2 Preprocessing

Due to the computational complexity of the PH calculation (typically, only small networks are considered for PH data analysis), the dataset size is reduced by taking the top 40% edges by mean NPMI 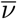 and mean NPMI rate, 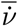.?

#### 3.2.1 Controls and Permutations

We use several different permutations as controls for our experiments, which are defined here. Given a dataset 𝒟_*w*_, we define the following permutations as functions 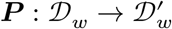 on the dataset: ***P***_*t*_, ***P***_*ν*_, and ***P***_*e*_.

***P***_*t*_ is a permutation of time wherein the ‘birth’ years for each edge is shuffled. This ablates the temporal and historical structure of the dataset. The node birth years are calculated and corrected given this permutation such that a node is assigned the year of the first edge which appears with that node as its boundary.

***P***_*ν*_ is a permutation of the *ν* values, wherein the entire array of *ν* over time for each edge is shuffled. This ablates the co-occurrence related structure of the dataset, while retaining the connectivity.

***P***_*e*_ is a permutation of the edges, by shuffling the *v*_*j*_ values for each (*v*_*i*_, *v*_*j*_) ∈ *E*. This ablates the connectivity of the dataset, while retaining the temporal information.

***P***_*y*_ is a permutation of the target labels in the link prediction task. This ablates the predictive power of any correlation between input and output of the dataset, while retaining any statistical abundance of certain classes.

### 3.3 Persistent Homology

See Section 8.3 for the basics on PH calculation.

In our experiments, we use the `Gudhi`^16–18^ library to compute the persistence using 2 different (related) filtration functions. Both of these methods were made to *approximate* the vineyard multiparameter persistence, by using a single parameter filtration on one of the parameters, and grading over another parameter.

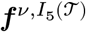 is a filtration over decreasing edge weights *ν* (increasing (1 − *ν*) ∈ [0, 2]), for each cumulative 5 years over the literature publication dates.

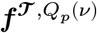 is a filtration over increasing time of the literature publication dates, for quantile bins of the edge weight *ν*. This filtration is more resolute over the time (individual years are used rather than 5 year bins), and less resolute over the (1 − *ν*) edge weighting parameter.

#### 3.3.1 Persistent Homology Centrality

For this work, we will define a persistent homology centrality measure (a centrality measure is *some* measure of the ‘importance’ of a vertex in a graph) based on the number of homology classes in which it is a member. In this work specifically, we choose to define it using only the cardinality of the *H*_2_ homological features in which the vertex participates:

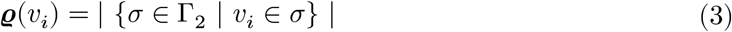

Where Γ_2_ is the set of representative *H*_2_ homological features in the graph, *σ* is the simplex that creates the *H*_2_ homological feature, and *v*_*i*_ is the vertex in question.

## 4 Results

### 4.1 Descriptive Statistics of the Dynamic Graphs

The baseline descriptive statistics of the co-occurrence network dataset 𝒟_2_ is shown in Figure 3, wherein the mean NPMI 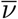 and mean NPMI rate 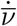 are shown for each edge (Figure 3A) along with the log-transformed document-level counts for nodes in Figure 3D. Individual edge information is shown in Figure 3B/C which displays the entire time series and evolution of the NPMI as the two entities either increase or decrease in their normalized co-occurrence over time. Similarly, the node information is shown in Figure 3E/F where the document counts for each node over time is displayed.

**Figure 1:**
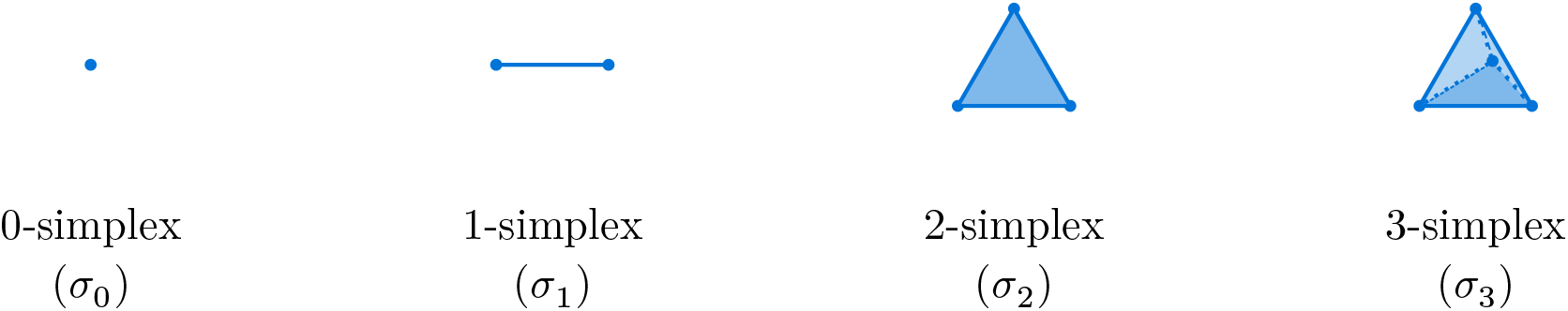
Simplices: Each *k*-simplex ***σ***_*k*_ spans a *k* dimensional space with *k* + 1 vertices.

**Figure 2:**
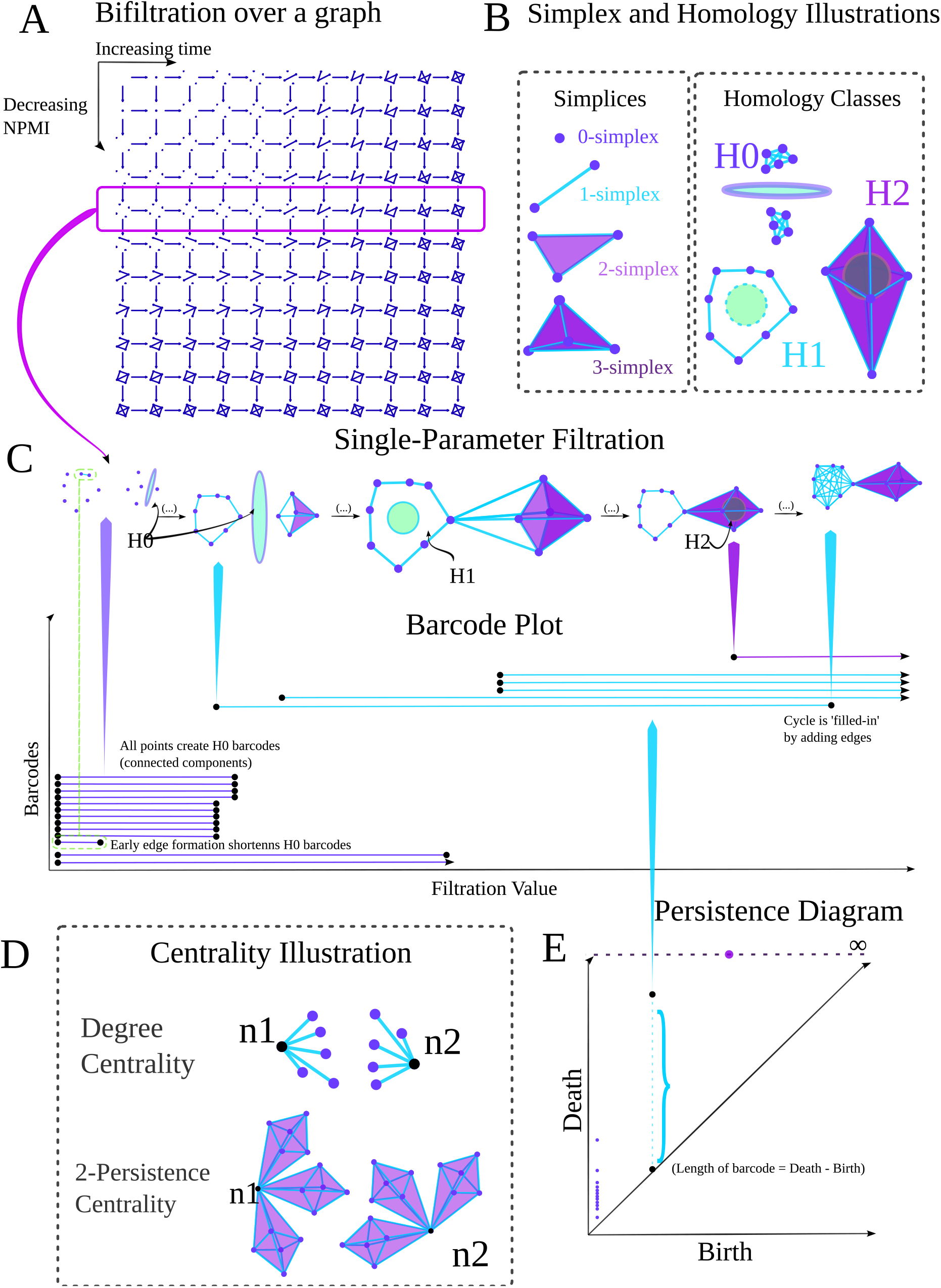
Overview of System: PH on the Pubmed network creates features for link prediction. **A**) *Bifiltration*: Grid of graph subsets along two parameters: time and (decreasing) NPMI. **B**) *Simplices & Homology*: Homology classes are often referred to as `k`_gap_s (this work), *cycles* (1D) or *voids* (2D). **C**) *Filtration & Barcodes*: A *single parameter filtration* creates a *barcode plot* which summarizes the persistence over the filtration. **E**) *Persistence Diagram*: A summary of the barcode plot in ℝ^2^ by a scatter plot. **D**) *Centrality Illustration*: The degree centrality is the # edges on a node. The *k*-persistence centrality is the # of *k*-homology generators on a node.

**Figure 3:**
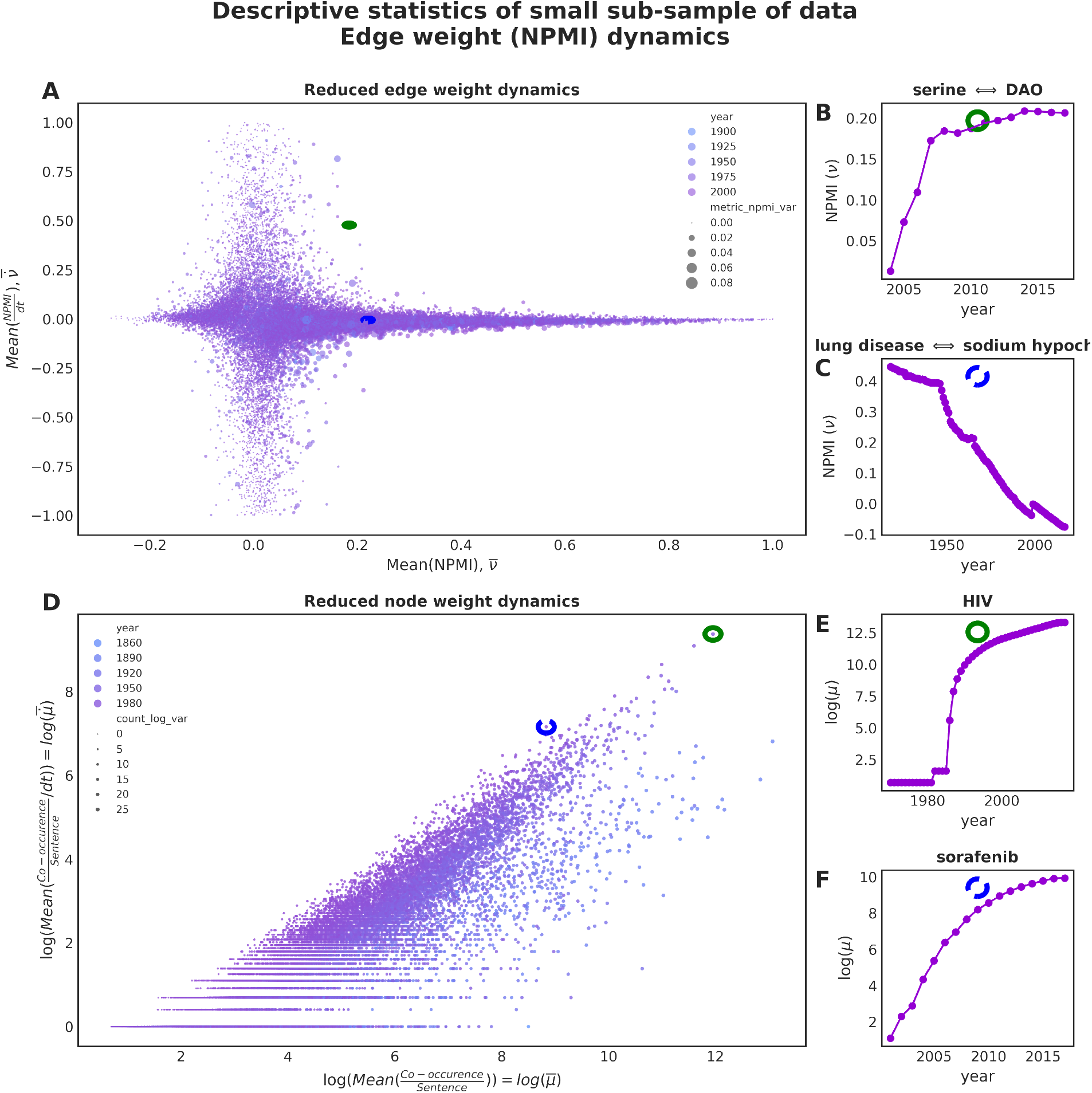
Descriptive statistics of dynamic datasets 𝒟_*w*_: General overview of the dynamic statistics of the network. **(A)** Aggregation of dynamic edge weight information for the network. The mean NPMI over all time points for each edge, i.e. 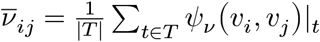 and mean NPMI *rate*, i.e. 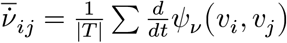. **(B**,**C)** Specific edges selected from (A), showing their full time series of NPMI. The inset green circle (B) or dashed blue circle (C) corresponds to the edge selected in (A). **(D)** Aggregation of dynamic node information for the network. The mean sentence-level co-occurrence counts over all time points for each node, i.e. 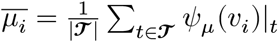 and mean sentence-level co-occurrence count *rate*, i.e. 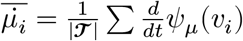. The document counts are log-transformed for visualization purposes. Plots from dataset 𝒟_2_, with a small sub-sampling of n=100,000

These specific nodes and edges were chosen to illustrate interesting points within the dataset by ranking in their 𝓁_2_ distance in the z-statistic space of (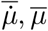, var(*μ*)) or (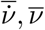, var(*ν*)).

### 4.2 Time-Graduated Persistence Diagrams

Applying a PH on a filtration ***f*** of increasing (1 − *ν*) to every 𝒢_*w,t*_ with time slices of every 5 years from 1960 to 2010, yields the persistence diagram shown in Figure 4. The persistence diagrams show that over time the median ‘lifetime’ over all `k`_gap_s (i.e. the median (birth − death) value) decreases, as the number of simplices increases (time). If the dynamic structure is ablated, then the simplices do not have the same connectivity opportunities early in the network, and therefore the median difference between NPMI values when simplices are connected is larger. This causes a large increase in the median lifetime of each homological feature.

**Figure 4:**
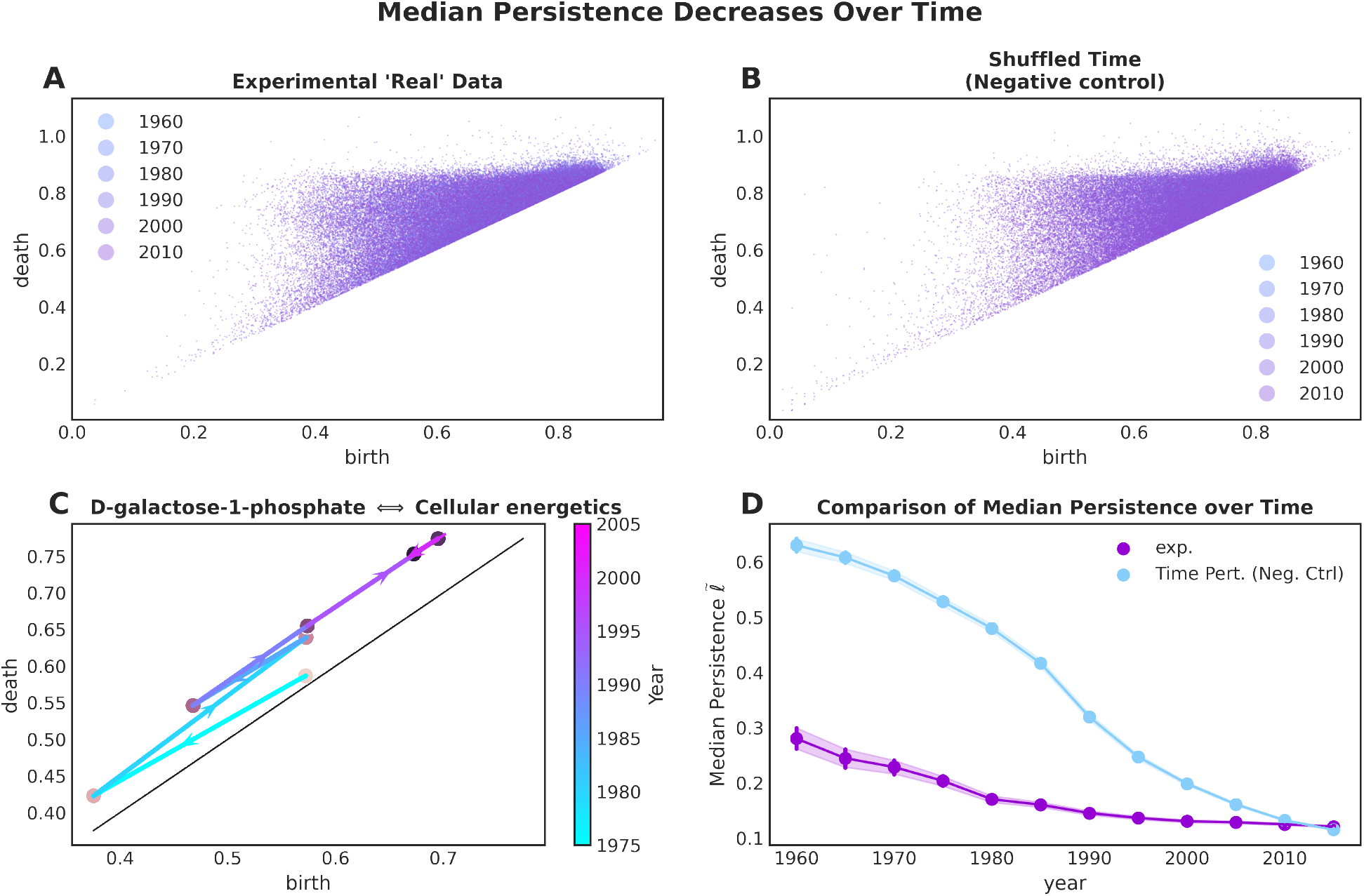
Time-graduated Persistence Diagrams: Scientists typically use the existing framework of knowledge and write about highly related concepts which are more tightly bound over time. Ablating the historical fidelity of the framework leads to a larger total persistence as the structure to form knowledge gaps from related phenomenon is not present. **(A)** Persistence diagrams for filtration over (1 − *ν*), graduated by time. **(B)** Persistence diagrams for time permutations ***P***_*t*_(𝒟_2_), with a filtration over(1 − *ν*), graduated by time. **(C)** Individual representatives of persistence generators can be tracked as *vines*. A representative homological generator is ‘born’ (i.e. “forms a knowledge gap”) when `D-galactose-1-phosphate` is introduced into the filtration. The filtration value (1 − *ν*) when `Cellular energetics` is added can be read from the ordinate for each year to determine how this homological feature changes over time. Homological features with infinite lifetimes are not shown. Plots from dataset 𝒟_2_ with a small sub-sampling of n=100,000

### 4.3 Link Prediction

The barcodes ℬ from the persistence calculations in the previous section are used to assign a single value ϱ_𝓂_ to each node within the graph. This value is determined by taking the cardinality of the membership of a node within each barcode above a certain homological dimension threshold 𝓂. This is similar to the Betti number in that it is an integer count of homological features; however, is is distinguished from a Betti number in that the Betti number ***b*** is a count of the number of generators (for any given birth simplex) of a given dimension, rather than tracked back to a node (0-simplex).

The result of this process is a centrality score, such that each node has a value assigned to it. This allows a comparison to one of the most common baselines for co-occurrence networks, the degree centrality. It is well known that degree centrality models co-occurrence graphs very well, and is a very efficient method to both produce networks that have similar properties to the co-occurrence network, and to predict future links within these networks. Therefore, casting the problem of link prediction as a classification problem, we show that the persistence-based centrality score ϱ_2_ outperforms the degree centrality in predicting future links in the co-occurrence network 𝒢_*w,t*_ in Figure 5 and Figure 6.

**Figure 5:**
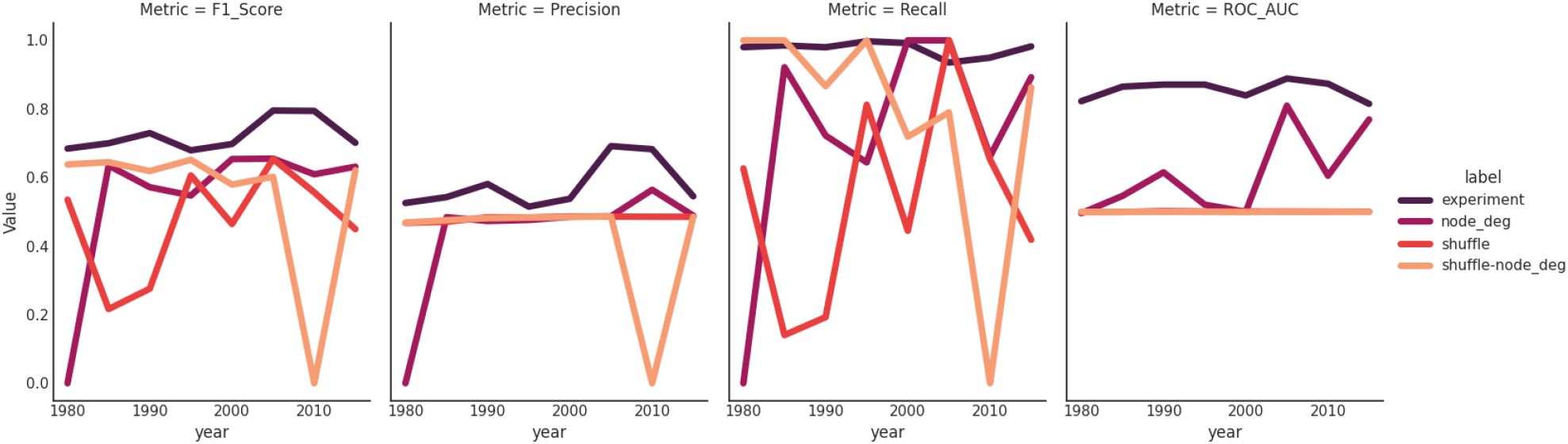
Link Prediction Performance: The node membership within ℬ_2_ is a good predictor of link existence within the network, improving upon every metric against the well-known simple baseline of degree centrality (preferential attachment). Perturbations on the target show ablation of performance.

**Figure 6:**
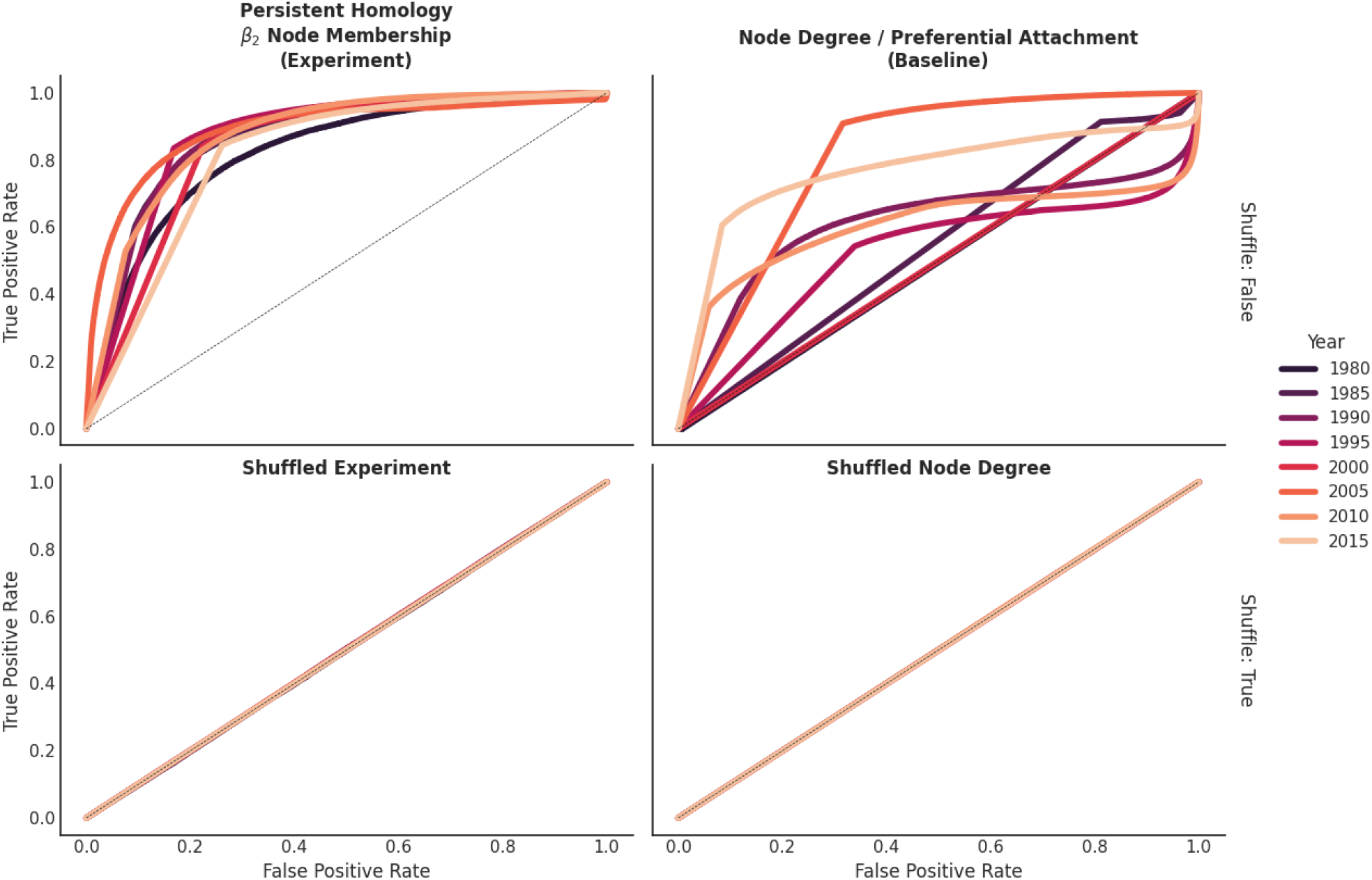
Receiver Operating Curves for the Link Prediction Task: The inability for ***P***_*y*_(𝒟_*w*_) to predict above random is more clearly shown in the ROC plots here. Our PH-based features are able to improve link prediction as a single feature in comparison to the baseline of degree centrality

**Figure 7:**
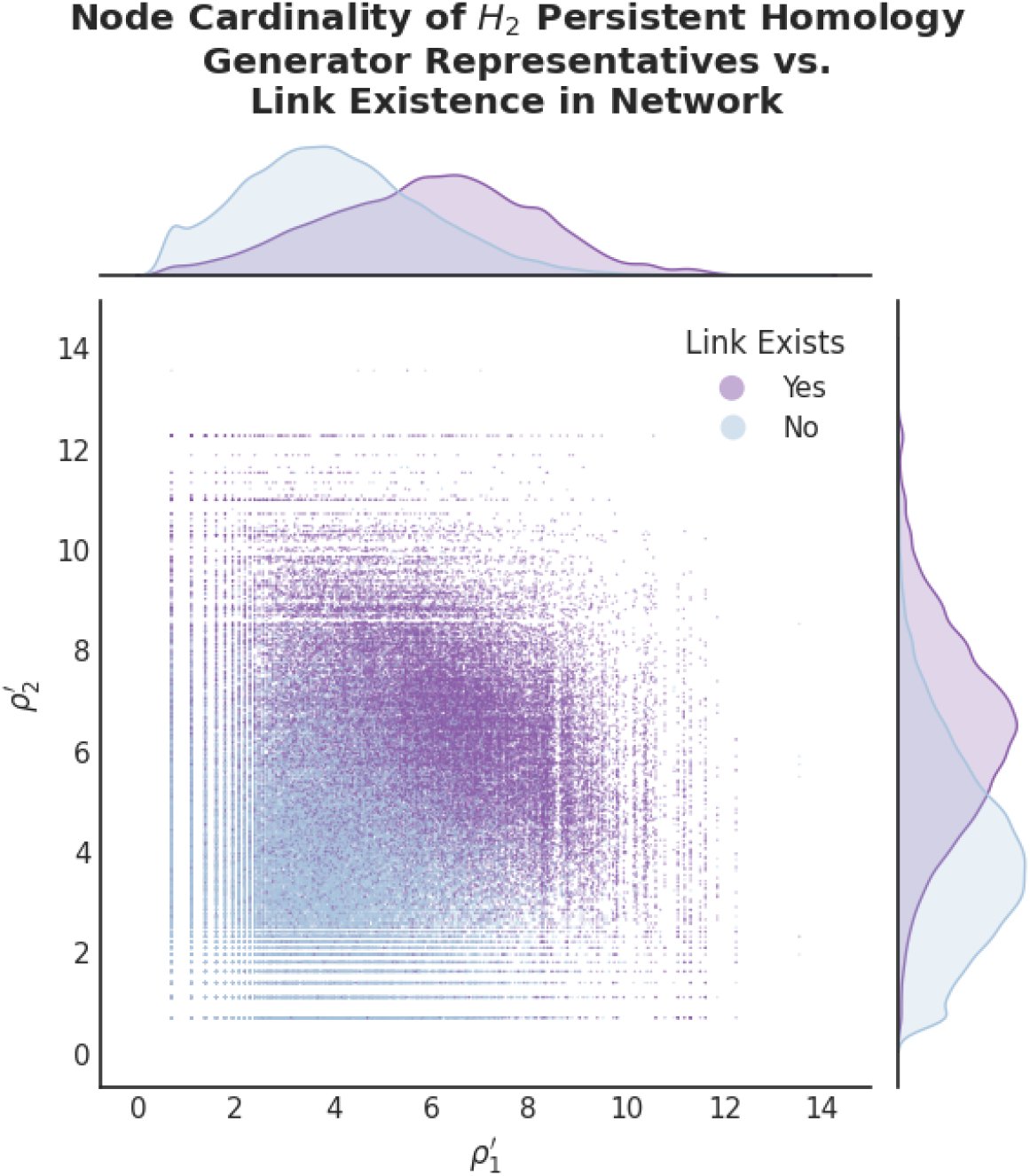
Link prediction data for PH centrality.

**Figure 8:**
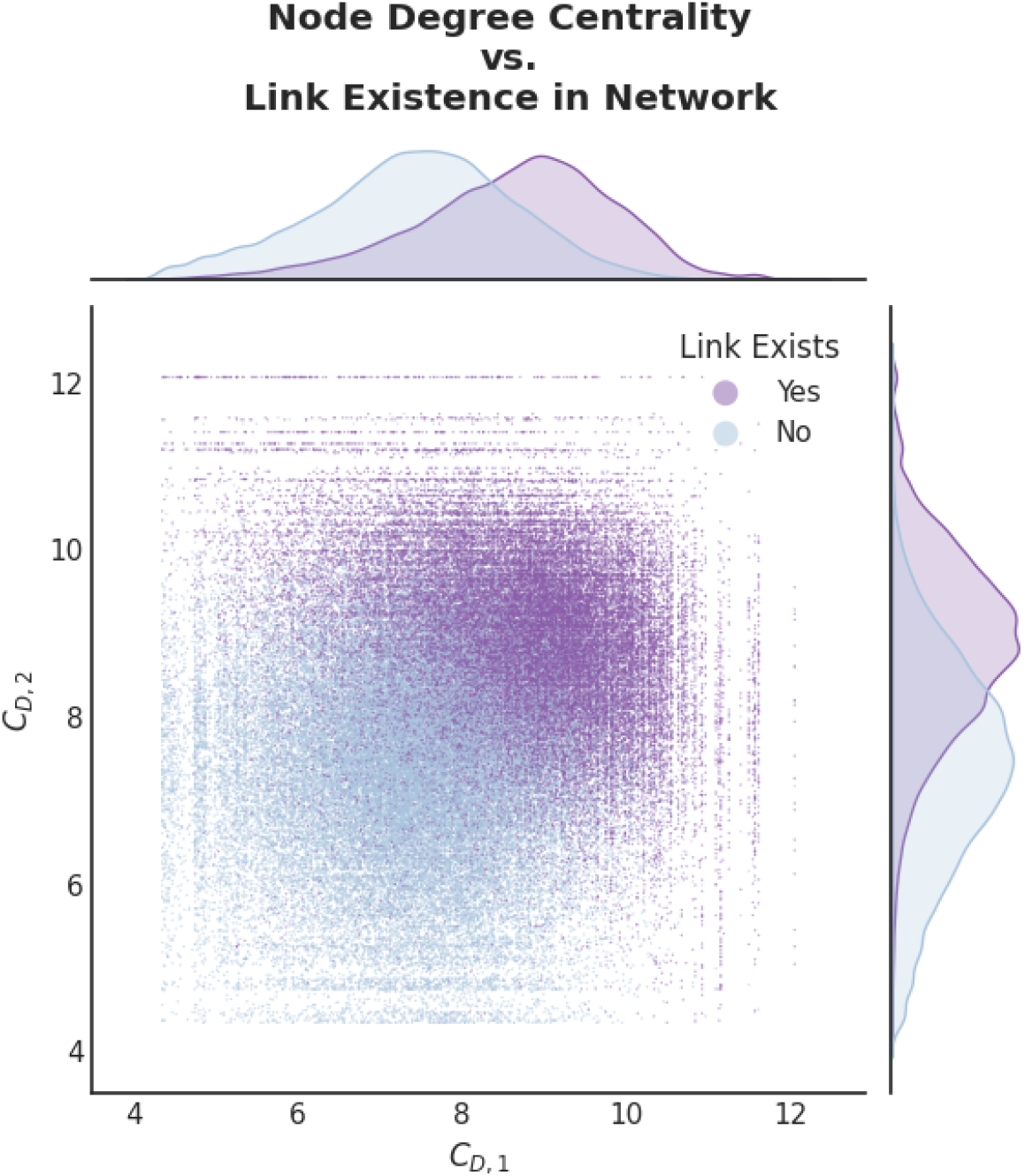
Link prediction data for degree centrality.

## 5 Discussion

The analysis provided suggests that algebraic topology - specifically persistent homology - can be a potent tool for uncovering dynamic structure in evolving networks generally, and is capable of discerning knowledge gaps within the scientific literature. This study underscores the intuitive positive correlation between answering a large and long-standing question, and the co-mention of the concepts that are key in describing the innovation (i.e. mutual information between concepts, normalized for abundance). Identifying these surges in NPMI of the simplex *ν* after filling a knowledge gap is particularly useful for researchers, as constructing a predictive model for identifying future scientific discoveries using this algebraic topology approach is a natural and operationalized process.

This research paves the way for the genesis of predictive tools in the field of literature based discovery, as well as scientific discovery more broadly. The identification and tracking of homological features in dynamic networks through other topological methods such as (more resolute) vineyards are also promising avenues for prospective further research.

This framing of new ‘ideas’, or sets of concepts, arising is intuitive, as if one thinks about a new method or biological entity or technology coming into the literature, it has to be mentioned within a framework of other concepts and described using the terms that define it in order to be understood. This sets the stage for the methodology used here, which simply operationalizes the process of identifying where the topological space is set up and primed for discovery to occur.

### 5.1 Mutual information dynamics

In this study, we have shown that the dynamic structure of the scientific literature can be partially captured within the mutual information between different biological entities in a co-occurrence network. This framework allows the detection of ‘knowledge gaps’ within this network via the application of persistent homology. In Figure 3, the landscape of established knowledge and trends within the literature can be easily identified. For instance, the ordinate 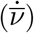 can be thought of as how *‘trending’* a set of entities is behaving across time, and the abscissa 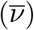 can be thought of as how *‘established’* a set of terms has been over time.

### 5.2 Persistent homology and representative cycles in the literature

#### A note on computational complexity

Due to the high density of edges in co-occurrence networks and the enormous computational complexity of persistent homology, the number of edges are pruned to the highest values of 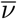 before expansion of the graph to include cliques of up to dimension *k* = 3. While a few advances have been made in the efficiency of persistent homology calculations on various fronts, these often have several constraints such as only operating on point clouds in ℝ^*d*^ (i.e. Vietoris-Rips Complexes have had considerable effort to improve their performance). There are also some improvements in Morse theory and Matroid filtrations^19^ that improve upon the standard historical performance from Delfinado, C. J. A. & Edelsbrunner, H. ^20^ (double exponential in the number of simplices, or linear in the number of simplices 𝒪(|𝒦|)) or methods using Smith-normal form.

We construct a “crude” vineyard (see Section 8.3.5) in Figure 4 to probe the dynamics of this system. The persistence diagram in Figure 4 shows that the process of scientific discovery leads to smaller ‘knowledge gap’ sizes over time, indicating that the questions scientists propose over time are more refined and specific. Small values for the total persistence indicate that the network is well-connected, and that the concepts written about in the articles are highly related. Conversely, shuffling the times for the edge information results in less connected networks wherein the pseudo-knowledge gaps are filled by ideas less related (as often the framework is not present in this control).

### 5.3 Predictive power of persistent homology in literature networks

As preferential attachment’s predictive power in link prediction within literature networks is a statement about probabilities (if two nodes both have a large percentage of the possible edges already incident upon them, it is likely that the edge in question will be one of them - particularly as the network radius shrinks), the predictive power of persistent homology features in link prediction is similar. In the case of 2-dimensional homological generators, nodes with large values will be similarly likely to share one of these edges.

## 6 Conclusion

This work has demonstrated that persistent homology can be used to identify and track knowledge gaps in scientific literature over time using methodologies similar to ‘large-step’ vineyards, or bi-filtrations for multi-parameter persistence. Our analysis reveals that an increase in the homology class 2 representative generators found by persistent homology is correlated with the probability that two concepts will be mentioned in the future. This type of link prediction improves upon past centrality-based link prediction methods, as it includes information about simplices and homological feature boundaries, unlike most other measures of centrality. This method therefore allows for a more performant model, while also maintaining explainability (unlike typical graph neural network models). These results suggest that persistent homology could be a useful tool for predicting scientific discovery, providing researchers with valuable insights into their areas of research and allowing methodologies for objective metrics of novelty when evaluating scientific literature and grant proposals.

## 7 Future Work

While our analysis has provided promising results, there are improvements to be addressed in future work. Firstly, continuous methods using Vietoris-Rips and multiparameter filtrations (such as complete resolution vineyards, or bifiltrations on density & distance) over language embedding vectors will uncover a more correct and resolute description of the underlying structure of the semantics of the data.

Another limitation of our study is that we only consider co-occurrence networks constructed from applying NER to the language within the abstracts directly, rather than using a directed graph structure constructed from extracting relations from the data. In future work, we plan to explore the use of different network representations and their impact on the results.

## 8 Appendix

### 8.1 Datasets

| $w$ | Dataset Name |
| --- | --- |
| 0 | landmark_nfkb-ademona |
| 1 | landmark_notch-cebpb |
| 2 | landmark_il17-mkp1 |
| 3 | landmark_nrf2-pancreatic_cancer |
| 4 | landmark_cxcl12-thyroid_cancer |
| 5 | swanson_migraine-magnesium |
| 6 | swanson_somatomedin_c-arginine |
| 7 | swanson_alzheimers-estrogen |
| 8 | swanson_alzheimers-indomethacin |
| 9 | swanson_schizophrenia-cipla2 |

### 8.2 Metrics

The point-wise mutual information (PMI) of a pair of entities, x and y is a measure of the information they provide about each other. Intuitively, it measures what is the probability of observing both x and y together, compared to the probability of observing x and y independently. It is defined below, as the logarithm of the probability of the pair divided by the product of the probabilities of the individual outcomes:

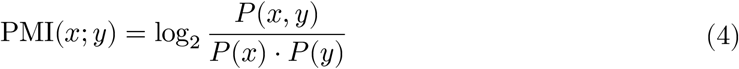

where *P* (*x, y*) is the probability of both x and y occurring together, and *P* (*x*) and *P* (*y*) are the probabilities of x and y occurring independently.

The *normalized* point-wise mutual information (NPMI) simply normalizes this to [−1, 1] by dividing by the negative of the logarithm of the probability of the occurrence of x and y together:

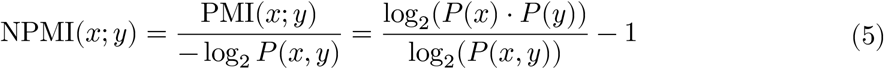

### 8.3 Persistent Homology

Persistent homology is a method from within the field of algebraic topology that specializes in analyzing the ‘shape’ of data by determining where and how long topological features persist. Constructing a persistence diagram consists of creating a *filtration* over the data, which is simply a set of inclusion maps - i.e. a sequence of nested subspaces of the data.

While there are many potential mathematical objects for which PH can be applied (such as the Vietoris-Rips complex for point cloud data), in this work we begin instead with a undirected graph structure.

#### 8.3.1 Simplicial Complexes and Filtrations

A *simplex σ* is simply an extension of the notion of nodes and edges to higher dimensions, for which the dimension of the simplex is defined by the dimension it spans, as seen in Figure 1.

These simplices have ‘faces’, which are conventionally denoted as *τ*, such that *τ* ⊂ *σ*. Similar to the way in which simplices extend the notion of nodes and edges these simplices can be ‘glued’ together to form a *simplicial complex* 𝒦, which is a generalization of a graph to higher dimensions (i.e. a type of ‘hypergraph’).

A *filtration* ***f*** assigns values to each of the simplices in a simplicial complex ***f*** : *σ* → ℝ, thereby ordering the simplices such that ***f***(*τ* ) ≤ ***f***(*σ*). Notice that this ordering allows the entire simplicial complex 𝒦 to be filtered, as each addition of a new simplex *σ* up to a threshold defines a simplicial complex ***f*** : 𝒦 → ℝ. This creates a *sequence*, or *ordered set of inclusion maps* of simplicial complexes, such that each complex is a subset of the next.

More formally, simplices *σ*, filtrations ***f***, and simplicial complexes 𝒦 have the following properties:

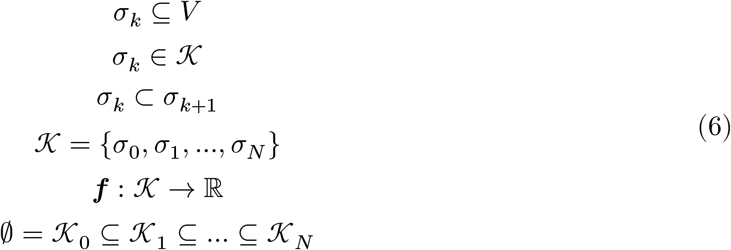

**Note:** Often a simplex tree constructed with `Gudhi` is also ‘expanded’ (`st.expansion(kmax)`) such that the simplicial complex can be ‘upgraded’ to including all cliques as simplices up to dimension *k*_max_, even if a *graph* 𝒢 (simplicial complex with *k* = 1) is provided, as is the case for this work.

#### 8.3.2 Chain Complexes, Boundary operators, and Homology

We define the boundary operator 𝜕_*k*_ as a linear map from the *k*-simplices *σ*_*k*_ to the (*k* − 1)- simplices *σ*_*k*−1_ (faces of *σ*). Additionally, we let *C*_*k*_(𝒦) be defined as an 𝔽_2_-vector space with basis given by the *k*-simplices of 𝒦 (where 𝔽_2_ is a field with two elements) such that:

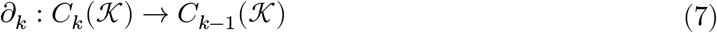

A *chain complex* is a sequence of chain groups connected by the boundary homomorphism:

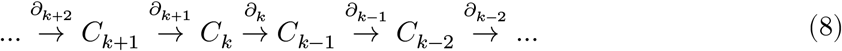

In other words, the boundary operator maps any *k*-simplex to it’s boundary.^21^

The homology groups *H*_*k*_(𝒦) are defined as the quotient of the kernel of the boundary operator for a *k*-simplex 𝜕_*k*_ by the image of the boundary operator for (*k* + 1)-simplices 𝜕_*k*+1_:

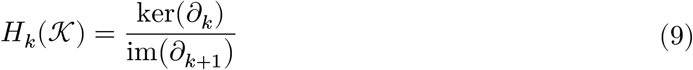

#### 8.3.3 Barcodes, Persistence Diagrams, and Representatives

Equipped with the homology groups over a simplicial complex *H*_*k*_(𝒦), we can now track the *barcodes* **ℬ** of the over the filtration. Barcodes are one of the main representations of persistent homology, which are multisets of intervals, where each interval is a tuple of (birth, death) filtration values, for which the persistence of the feature (death - birth) is often referred to as the ‘lifetime’.

To reiterate, each homology class (*H*_0_, *H*_1_, …, 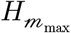) has a set of intervals (i.e. the *multiset*), where each interval (*b, d*) is the filtration value which creates or destroys a homological feature.

The barcodes **ℬ** are then used to construct the *persistence diagram* 𝒫, which is a multiset of points in the plane ℝ^2^. Each point in the persistence diagram represents a homology class together with its birth and death times.

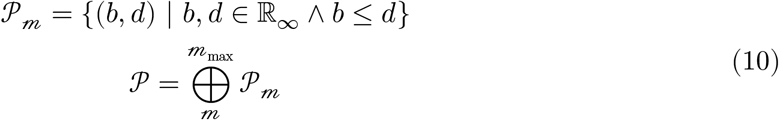

where ℝ_∞_ is the extended real number line, such that ℝ_∞_ = ℝ ∪ {∞}, as some homology classes may persist indefinitely.

Further, the *representatives* Γ of the persistent homology generators are the simplices that are responsible for the creation and destruction of the homology classes. These representatives can tie the points in the persistence diagram back to the original data if the original simplex was ‘labelled’ in the original data.

#### 8.3.4 Betti Numbers and Betti Curves

Betti numbers ***b***_*k*_ are a set of numbers that represent the rank of the homology groups *H*_*k*_ of a simplicial complex 𝒦. The *k*-th Betti number ***b***_*k*_ is defined here:

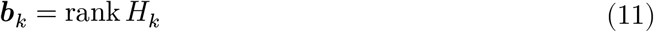

The first few Betti numbers have interpretable explanations such as ***b***_0_ being the number of connected components, ***b***_1_ being the number of loops, and ***b***_2_ being the number of voids in the space.

A related representation of the Betti numbers is the *Betti curve* ***β***, which is a multiset that tracks the Betti numbers over a filtration. The Betti curve can also be thought of as the ‘density’ of the barcodes **ℬ** in a barcode plot.

#### 8.3.5 Vineyards

A vineyard is a type of multi-parameter filtration for persistent homology that utilizes *time* as a parameter, as well as another parameter, such as Euclidean distance in the Vietoris-Rips complex. The vineyard is a plot of the persistence diagrams with a *third* dimension of time, which creates a plot that resembles a ‘vineyard’, in that several 1D lines (generators tracked over the *applicate*, or ‘z-axis’) curve through this 3rd dimension.

## 9 Glossary

Symbol: Definition
𝒟: Collection of datasets
*w*: Dataset indexer
𝒟_0_: First dataset (i.e. collection of `.tsv`s for `landmark_nfkb-ademona`)
𝒢_*w,t*_: Graph of dataset *w* at time *t*
*I*_Δ_(*S*): Interval function acting upon a set *S* to create a set of intervals with step Δ
*Q*_*p*_(*S*): Quantile function acting upon a set *S* to create a *set* of quantiles with *p* elements, i.e. *Q*_10(*S*)_ is all 10 deciles of *S*
*𝒯*: Set of *all* times provided within a dataset
*I*_Δ_(𝒯): Set of time interval *slices* created from a dataset. ∴ *I*_Δ_(𝒯) = {[*t*_*i*_, *t*_*i*+Δ_] | *t*_*i*_, *t*_*i*+Δ_ ∈ ***𝒯*** ∧ *t*_*i*_ < *t*_*i*+Δ_}
*T*: Number of time interval slices created from a dataset. *T* = |*I*_Δ_(𝒯)|
*t*_0_, *t*_*f*_: Initial and final time within the interval slice set *I*_Δ_(𝒯).
*V*: Set of vertices in a graph *G*
*v*_*i*_: Single vertex indexed as *i*. An element in *V* . Not to be confused with *ν*_*i*_ (the NPMI of *v*_*i*_)
*E*: Set of edges in a graph *G*
*e*_*ij*_: Single edge (*v*_*i*_, *v*_*j*_) in a graph *G*
Φ_ν_: The set of functions which map vertices to a value (such as *φ*_*μ*_ ∈ Φ_ν_ where *φ*_*μ*_ : *v*_*i*_ → *μ*)
Φ_*e*_: The set of functions which map edges to a value (such as *φ*_*ν*_ ∈ Φ_*e*_ where *φ*_*ν*_ : (*v*_*i*_, *v*_*j*_) → *ν*)
*ν*: Normalized point-wise mutual information (NPMI) between vertices
*φ*_*ν*_: NPMI function which operates on a pair of vertices (*v*_*i*_, *v*_*j*_)
*φ*_*μ*_: Document co-occurrence (co-mention) function. This is the sum of all co-occurrences of this term in the dataset at the sentence level, and is referred to as ‘count’ in the original table’s column
*φ*_#_: Document-level count function. This is the sum of all occurrences of this term in the dataset at the document level, and is referred to as ‘doc-count’ in the original table’s column
*σ*_*k*_: Simplex of dimension *k*
*τ*: Face of a simplex *σ*, ∴ *τ* ⊂ *σ*
𝒦: Set of simplicial complexes for a dataset over a time range 𝒯
𝒦: Simplicial complex. Constructed from ‘gluing’ a set of simplices together, or ‘expanding’ a graph to include higher dimensional simplices (e.g. triangles) from any cliques present within it
*f*: Filtration function to operate on simplicial complexes. ***f*** : 𝒦 → ℝ
*f*(*σ*): Filtration value of a simplex *σ*
𝒦_0_, 𝒦_*f*_: Simplicial complex at first index, or final simplicial complex index *f*
*k*_max_: Maximum dimension of simplices in **𝒦**, i.e. |*σ*_*k*_| ≤ *k*_max_∀*σ* ∈ **𝒦**
𝓂_max_: Maximum dimension of homology class to be calculated for persistent homology
𝜕_*k*_: Boundary operator
*C*_*k*_: Chain group
*H*_*k*_: Homology group
*b*_*k*_: Betti number
*β*: Multiset of Betti curves over a filtration for each *k*
*β*_*k*_: Betti curve for a specific *k*
ℬ: Collection of barcodes
*𝒫*: Collection of Persistence Diagrams
𝒫: Persistence Diagram
Γ: Representatives of persistent homology generators
ϱ_*k*_: Representatives of homology class k generators
`k`_gap_: “Knowledge gap” - This is a loose synonym for a homology class *H*_*k*_, or even a specific representative cycle Γ

